# Specific Virus-Host Genome Interactions Revealed by Tethered Chromosome Conformation Capture

**DOI:** 10.1101/142604

**Authors:** Haochen Li, Reza Kalhor, Bing Li, Trent Su, Arnold J. Berk, Siavash K. Kurdistani, Frank Alber, Lin Chen

## Abstract

Viruses have evolved a variety of mechanisms to interact with host cells for their adaptive benefits, including subverting host immune responses and hijacking host DNA replication/transcription machineries [1–3]. Although interactions between viral and host proteins have been studied extensively, little is known about how the vial genome may interact with the host genome and how such interactions could affect the activities of both the virus and the host cell. Since the three-dimensional organization of a genome can have significant impact on genomic activities such as transcription and replication, we hypothesize that such structure-based regulation of genomic functions also applies to viral genomes depending on their association with host genomic regions and their spatial locations inside the nucleus. Here, we used Tethered Chromosome Conformation Capture (TCC) to investigate viral-host genome interactions between the adenovirus and human lung fibroblast cells. We found viral-host genome interactions were enriched in certain active chromatin regions and chromatin domains marked by H3K27me3. The contacts by viral DNA seems to impact the structure and function of the host genome, leading to remodeling of the fibroblast epigenome. Our study represents the first comprehensive analysis of viral-host interactions at the genome structure level, revealing unexpectedly specific virus-host genome interactions. The non-random nature of such interactions indicates a deliberate but poorly understood mechanism for targeting of host DNA by foreign genomes.

## Background

Upon infection, viruses deliver viral proteins and genetic materials into host cells either embracing or confronting the host cellular machineries. A variety of virally evolved adaptive mechanisms lead to dynamic virus-host interactions at every stage of the viral life cycle—all intended to increase the likelihood of generating progeny virions. A key aspect of viral regulation is the transcription of viral genes and the replication of viral genomes. Many RNA and DNA viruses introduce DNA into the host cell nucleus. Retrovirus and lentivirus RNA genomes also enter the nucleus as double stranded DNA, which is synthesized by virion-associated reverse transcriptase in cytoplasm. The genome of DNA viruses is packaged at very high molecular density with specialized viral packaging proteins to form a capsid virion particle. Upon entering the cell nucleus, virion particles interact with host activating factors in decondensing the incoming viral DNA to initiate viral transcription and replication or establish a stable intermediate for latent infection or long-term integration [4,5].

After nuclear entry, the viral genomes distribute inside the nucleus through a non-random but poorly understood process. For DNA viruses, fluorescence in situ hybridization (FISH) imaging analyses show nuclear deposition of viral DNA and subsequent viral gene transcription at the periphery of Nuclear Domain 10’s (ND10s), which are also known as PML nuclear bodies (NBs) and PML oncogenic domains (PODs) [6,7]. ND10 is a highly organized nuclear structure accumulated with interferon-upregulated proteins, implicating ND10s as sites of a nuclear defense mechanism. ND10 subcompartments often reside adjacent to nuclear speckles, a nuclear domain enriched in pre-mRNA splicing factors. DNA viruses initiate their transcription at ND10s juxtaposed to nuclear speckles. After the expression of viral immediate early gene products, ND10s subsequently become dispersed. The dynamic co-localizations of DNA virus genomes and host nuclear domains at defined stages of viral life cycle strongly suggest a functional role of viral-host genomic interactions. However, because of the limited resolution of FISH and its low throughput, the molecular and chromatin compositions of these nuclear domains and their interactions with viral DNA are not fully characterized. Little is known about whether the virus genome associates with specific host chromosomal regions.

Extensive studies suggest that nuclear organization, including the proximity of genomic loci and their positions inside the nucleus, play important roles in regulating transcription and DNA replication [8–10]. In addition to early image-based analyses, recently developed chromosome conformation capture (3C) techniques are able to capture chromosomal interactions at fine resolution and in high-throughput (Hi-C) when coupled with next generation sequencing [11,12]. These studies reveal a hierarchical organization of chromosome structures—including contact domains [13], topological associated domains [14], macro-domains [15], and chromatin compartments A/B [12] —that can be functionally related to gene expression, chromatin modifications, DNA replication timing, etc. These studies raise the possibility that viruses may have to adapt to or alter the host nuclear environment and genome structure to regulate the viral life cycle. To reveal whether virus-host genome interactions are specific, we used a 3C/Hi-C based approach to investigate the viral host genome interactions. However, solution-based 3C methods such as Hi-C suffer from high background noise [15–18], such that rare DNA interactions, including interchromosomal contacts as well as virus-host genome interactions to be studied here, may be difficult to detect. Alternative approaches, such as the solid phase chromosome conformation capture technique, also known as TCC [15], and the recently developed *in situ* Hi-C and single-cell Hi-C [13,19], minimize the noise levels originating from random DNA ligations and could be adopted to investigate virus-host genome interactions. In this study, we used TCC and the adenovirus/fibroblast model to explore virus-host genome interactions and its potential functional implications.

The adenovirus is a non-enveloped virus containing a linear double stranded DNA genome. It is the first human virus that was found to cause tumors in hamster [20]. This tumor promoting activity led to extensive studies of the molecular and cell biology of adenovirus infection [21]. Among the human adenoviruses, serotype 5 (Ad5) is an extensively characterized strain, which has a ~36 kb linear dsDNA genome and encodes ~39 viral genes [22].

Here we identified genome-wide adenovirus-host genome interactions and observed that the interactions are not randomly distributed across the host genome. Host-viral genome interactions preferentially occur in specific regions of the host genome that are associated with open and early replicating chromatin. At the same time, these regions are also enriched with specific histone modifications, such as H3K27me3, which is not normally associated with active chromatin but is critical in regulating cell differentiation [23]. Interestingly, the viral DNA attachment sites coincide with locations undergoing early host epigenome remodeling upon viral infection. We found a correlation between the observed frequencies of a host-viral DNA interaction and the extent of acetylation of H3K18 and H3K9 (in the presence of RB and p300), which are part of the observed epigenetic remodeling of the host genome upon adenoviral infection [24–27]. This observation raises the possibility that the locations of viral DNA in the host genome may be mechanistically linked to the dramatic epigenetic changes in the host genome upon viral infection.

## Results and Discussions

### Genome-wide interactions between adenovirus and host genome

We used TCC to map genome-wide chromatin interactions between the adenovirus and host genomes. We used the small E1A (e1a) adenovirus 5 construct (*dl1500*) that mainly expresses the small e1a oncoprotein [28]. The ‘early region 1a’ (E1A) of the human Ad5 genome generates two splice variants, the smaller of which (i.e., small e1a) is responsible for activating early viral genes’ expression and the virus-induced host cell transformation. The e1a oncoprotein can drive contact-inhibited human fibroblast cells into S phase in part by displacing the RB cell cycle checkpoint protein from the E2F master transcription factors [29] without causing host nucleus destruction in the lytic infection cycle. Extensive studies have demonstrated that e1a is crucial for remodeling the host epigenome leading to oncogenic transformation of the cell [24–26,29]. Thus, the *dl1500* construct serves as a good model for our studies. TCC experiments were performed on human primary lung fibroblasts (IMR90) with and without *dl1500* adenovirus infection for different durations at 6 and 24 hours post infection (p.i.). For the two datasets with virus infection, we preprocessed (Methods) the pair-end sequencing reads and aligned them to both human (hg19) and adenovirus 5 (Ad5) genomes. Pair-end reads aligned to hg19 on one end and to Ad5 on the other end were described as virus-host genome interactions (VGIs). 156,666 VGIs and 460,988 VGIs were identified at 6 and 24 hours p.i., respectively. Pair-end reads with both ends aligned to hg19 are host chromosomal interactions (CIs). For intrachromosomal interactions only pair-end reads with end reads separated by a sequence distance of at least 20 kb were considered. The absolute numbers of reads for each interaction type (CIs and VGIs) in all four datasets are shown in Table S1. VGIs consist of 0.12% and 0.43% of all detected interactions in the 6 and 24 hours p.i. data sets, respectively (Figure 1a).

**Figure 1.**
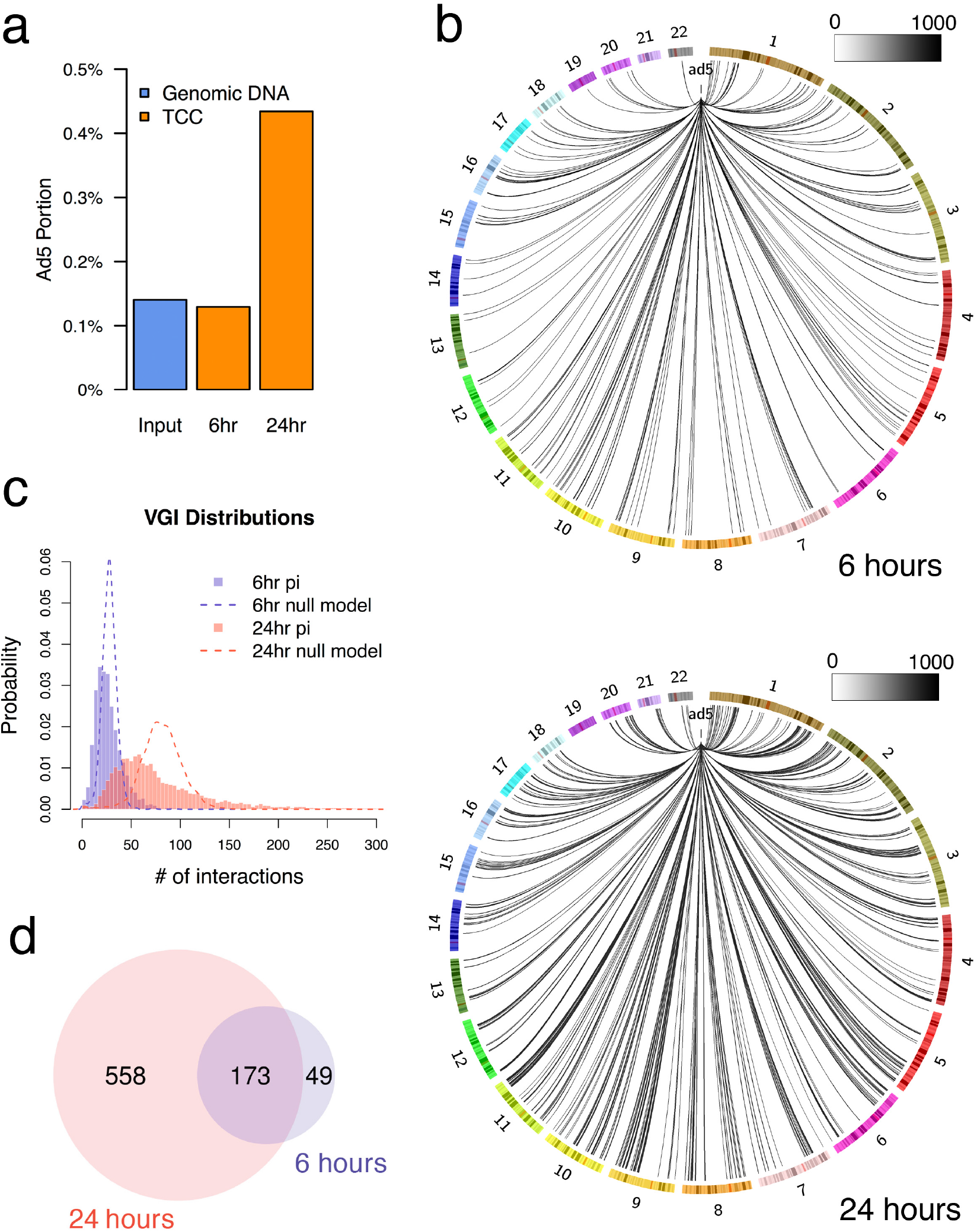
VGI distributions and enriched regions on host genome. a) Portion of TCC contacts containing adenovirus 5 (Ad5) DNA (including VGIs and viral-viral genome interactions). Input represents the portion of adenovirus DNA among the total DNA material in the nucleus, which is calculated by Size_Ad5*MOI (200 for the two virus infection TCC experiments in this study) / (Size_diploid_hg19+Size_Ad5*MOI). b) VGI contact frequency distributions on different chromosomes at 6 hours and 24 hours pi. VGI contact frequency vectors were normalized by the ICE bias vectors (Methods). Only statistically VGI enriched host genomic regions were plotted with lines (line gray scale represents VGI contact frequency) on the circos plots. c) Distributions of VGI observed contacts in the 500kb resolution bins at 6 hours and 24 hours pi. Each dash line represents the multinomial null model distributions assuming VGIs are randomly distributed across the host genome. d) Venn plot of VGI enriched bins identified at different time of infection using the null models given FDR < 10^−4^.

The amount of adenovirus DNA in the nucleus of an infected cell can be calculated from the size of the adenovirus 5 genome (~36 kb) multiplied by the multiplicity of infection (MOI), which is the ratio of the infectious adenovirus particles to the number of targeted fibroblast cells. An MOI of 200 was used for all virus infection experiments to ensure near complete infection of all cells in culture. The total amount of DNA consists of the diploid female human genome (IMR90) and the viral DNA. The expected DNA capturing rate, defined as the ratio of the viral DNA to the total amount of nuclear DNA, is 0.14%, which is similar in value to the total fraction of VGI among all detected DNA interactions in the 6 hours p.i. TCC dataset (Figure 1a). This value indicates the TCC method is robust in capturing virus-host genome interactions.

To analyze the genome-wide distribution of VGIs, the host genome was tiled into binning windows at different resolutions. Given the current sequencing depth, a resolution of 500 kb binning windows was chosen as the optimal resolution to represent VGIs at the given sequencing depth (Methods and Figure S1). Chromosomal interactions (CIs) were also binned to 500 kb windows to build the Observed genome-wide raw Contact count (OC) matrices.

For 3C-based experiments, the number of contacts observed for different genomic regions may be biased by differences in DNA sequence and chromatin features, such as restriction enzyme digestion efficiency, sequence mappability, GC content, etc [30–32]. Based on the “equal visibility” assumption, the genome-wide OC matrices of the four datasets were normalized by balancing the marginal sum of the matrices through the iterative correction (ICE) method to obtain the contact frequency (CF) matrices [33] (See Methods for the details of data processing). To accurately account for the VGI contact frequency, the observed raw counts of VGI were normalized by the bias vector generated from ICE normalization method (Methods). VGI contact frequencies at 6 and 24 hours p.i. were therefore represented by two contact profile vectors with dimension of 5,689 bins spanning the entire hg19 genome. The Pearson’s correlation coefficient between the two vectors is 0.768 indicating that VGI positions are consistent between the two samples and are not randomly distributed. Genome-wide VGIs together with their contact frequencies are displayed in figure 1b, which shows the non-uniform distribution of VGI’s in the human genome for both datasets at 6 and 24 hours p.i.

### Identification of viral DNA enriched regions on the host genome

The enrichment of VGIs at a genomic region was determined by hypothesis testing with a null model assumption of multinomial distribution (Figure 1c; Methods). As results, 222 and 731 bins were identified as viral DNA enriched bins for the 6 and 24 hours p.i. samples, respectively. 173 (79%) of the VGI enriched bins in the 6 hours p.i. data set were also found in the 24 hours p.i. sample (Figure 1d). Since viral DNA does not replicate at early infection stages [24], the 222 viral DNA enriched bins identified at 6 hours p.i. are likely the primary genomic locations where adenovirus DNA interacts with the host genome as it is delivered into the host cell nucleus.

### Viral DNA enriched regions co-localize in 3D space prior to infection

Previous FISH imaging analyses showed that the adenovirus genome is deposited to specific nuclear domains (ND10s) at the early stages of viral infection [7]; and that ND10s segregated equally and had symmetric positions in daughter nuclei after cell division [34], which indicates viral DNA genome deposition follows certain chromosomal bases. Moreover, ND10s are frequently situated next to DNA replication foci and specific gene loci [6]. These observations suggest that the viral genome prefers certain larger host chromosomal locations or subcompartments within the nucleus.

To investigate the existence of large chromosomal subcompartments involved in viral-host genome interactions, we tested if the 222 chromatin regions enriched with VGIs at the 6 hours p.i. are clustered in 3D space before the viral infection. The 222 bins can form up to 1507 bin pairs of intrachromosomal interactions in the 6 hours mock-infection (m.i.) TCC dataset (Figure 2a). Within a chromosome, genomic regions generally tend to interact more frequently when they are separated by shorter sequence distances. Therefore, we performed a sequence distance restricted sampling test to rule out bias with respect to sequence proximity on the interaction frequency. For each intrachromosomal pair of the 1507 VGI regions, a pair of bins is randomly selected from the same chromosome with identical sequence distance separation to the VGI pair. The distance restricted sampling procedure was repeated for 10,000 times, generating 10,000 random bin pairs of intrachromosomal interactions. In this analysis VGI bins had a statistically significantly higher probability (p-value less than 10^−4^) to colocalize with each other in comparison to other bins with the same sequence separation, as is evident from the distributions of pairwise contact frequencies for VGI regions and random control (Figure 2b). Intrachromosomal long-range interactions define chromosome sub compartments, such as compartment A and compartment B regions. Chromosome regions within each compartment tend to co-localize with other regions in the same compartment [12]. On one hand, out of the 5,689 500-kb bins in the genome, 2483 (43.6%) are classified as compartment A regions in the 6 hours p.i. TCC dataset. On the other hand, out of the 222 500kb bins enriched with VGIs at 6 hours p.i., 148 (66.7%) are classified as compartment A regions. Therefore, the observed VGI colocation could be a reflection of its compartment A preference. To investigate if the VGI colocation is influenced by compartment A/B segregation, we repeated the distance restricted sampling test for the 148 VGI bins in compartment A in comparison to all the 2483 bins in compartment A. Also in this analysis, the p-value was less than 0.005, which indicates that viral genome colocation is not solely driven by its compartment A preference.

**Figure 2.**
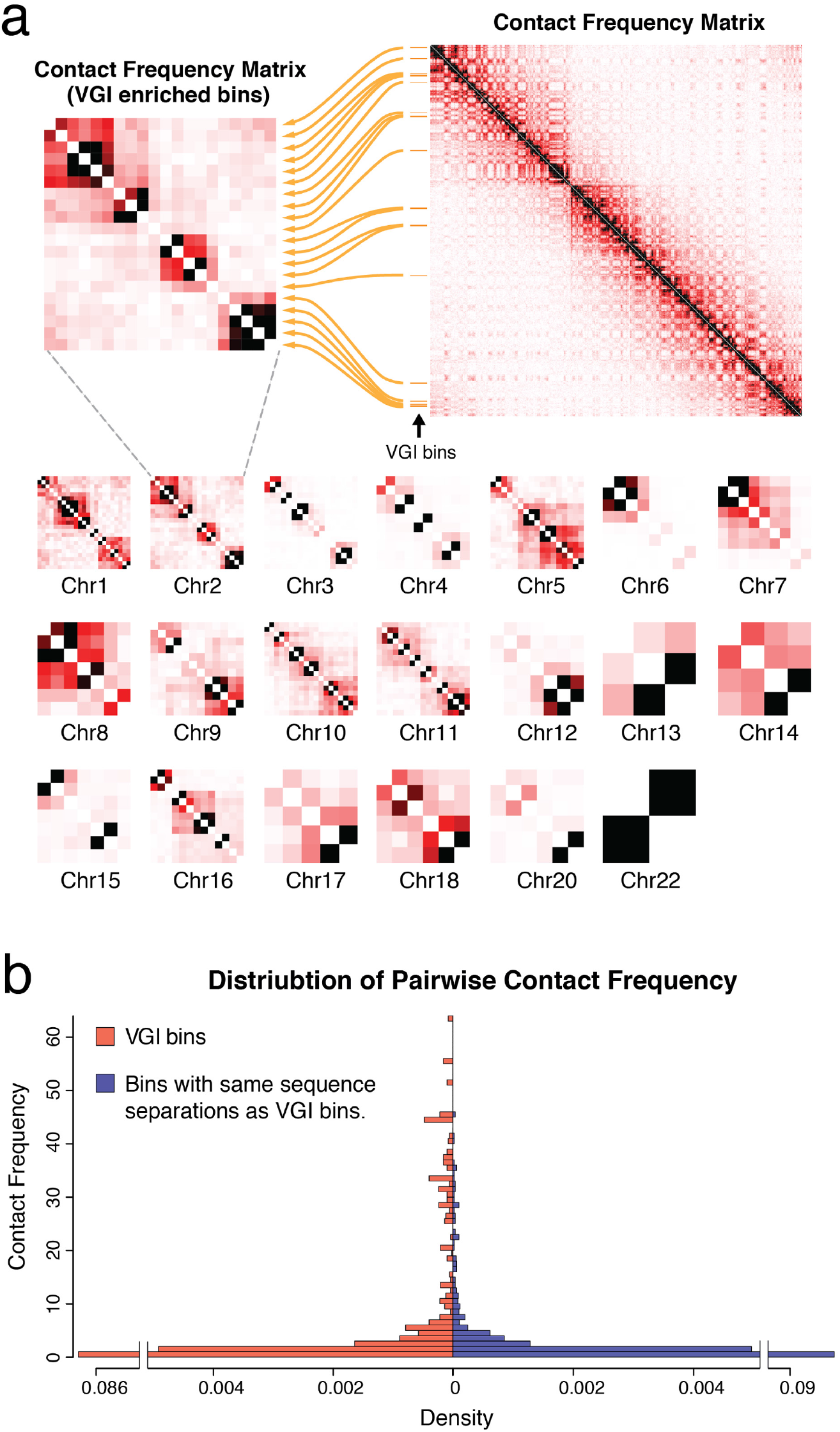
Spatial co-localization of VGI enriched chromatin regions. a) VGI enriched bins (6 hours pi) are selected from the 6 hours mock-infection TCC chromosomal contact frequency matrix. The selected sub-matrix shows a number of spatial clusters for different chromosomes. Sub-matrix for chromosome 21 is not shown in the figure, since only one VGI enriched bin is identified in the chromosome. Similarly, the two VGIs enriched bins for chromosome 22 are consecutive located on the chromosome. Therefore, contact frequency between these two bins is substantially high. b) Distribution of intrachromosomal contact frequencies of the selected sub-matrices is shown on the left side (in red). Distribution of contact frequencies generated by the distance restricted sampling method is shown on right side (in blue).

Interestingly, our analysis indicates that VGI bins can be clustered according to their contact frequency profiles into 2 to 3 larger groups (i.e. subcompartments) per chromosome, so that VGI regions within each domain showed substantially increased interaction frequencies in comparison to interactions to VGI regions in another domain. These spatial subcompartments are visually evident when plotting the contact frequency heatmaps formed by only VGI bins (Figure 2a). These subcompartments could be candidates for nuclear domains associated with early viral infection, such as the previously identified ND10s [7].

### Viral DNA preferentially interacts with active host chromatin regions and chromatin regions marked by histone modification H3K27me3

The genome-wide chromosome conformation capture data of IMR90 is able to classify the host genome into at least two distinct compartments, A and B, based on the TCC interaction patterns [12,15]. In addition to the spatial distinction between different compartments, compartment A regions are also characterized by active chromatin features including an early replication timing profile, while compartment B regions are recognized as “inactive” regions [12,15]. Figure 3a shows the assignment of compartment A and B for chromosome 2. For the entire genome, VGI frequencies for regions of compartment A were significantly higher than those in chromatin regions of compartment B (with student’s t-test p-value less than 10^−15^) (Figure 3b). Among the 222 VGI enriched bins at 6 hours p.i., 148 (66.7%) belong to compartment A, while for the whole genome 2483 (43.6%) out of 5689 bins are part of compartment A. VGI frequency distributions for regions in compartments A and B in all chromosomes are detailed in Table S2.

**Figure 3.**
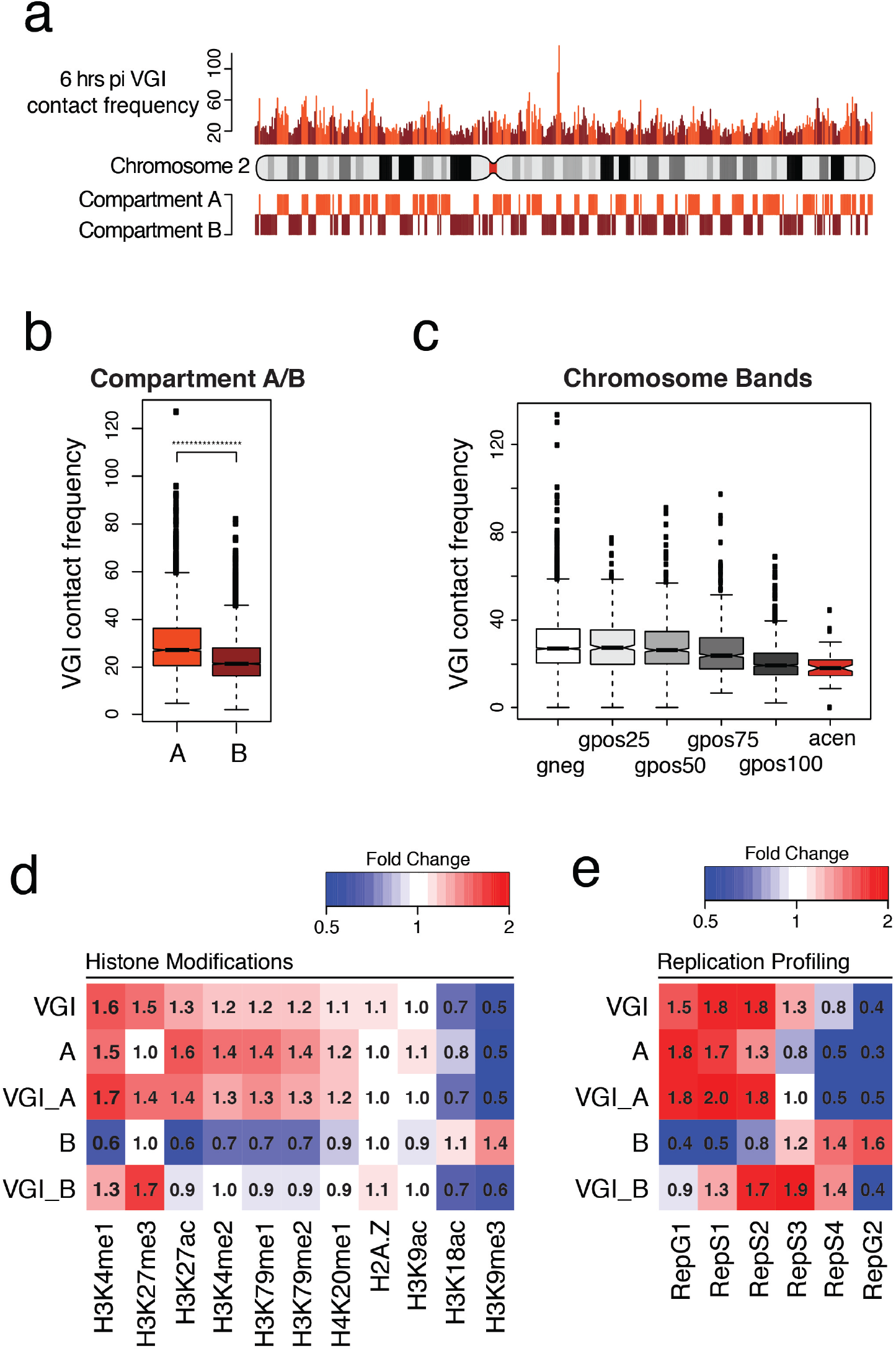
Adenovirus genome interacts with active chromatin regions and early replicated genomic regions. a) Chromosome 2 cytogenetic features and compartment A/B assignments are shown within the schematic chromosome. 6 hours pi VGI contact frequency profile of chromosome 2 is shown on top and colored according to the A/B compartment membership of the corresponding chromatin region. Cytogenetic features, compartment A/B assignment, and VGI contact profile are all shown at 500kb resolutions. b) Box plot of VGI contact frequencies for chromatin of compartment A and B bins at 6 hours pi. c) Box plot of VGI contact frequencies for chromatin regions of different cytogenetic categories at 6 hours pi. d) Heatmap of different histone modification markers and replication timing signals for different categories of genomic regions, including, all VGI enriched regions, all compartment A regions, VGI enriched compartment A regions, all compartment B regions, and VGI compartment B regions. Strengths of signals for the heatmap entries are represented by fold change compared to the genome-wide average signals. H3K27ac data is from Ferrari et al. 2014 [27]. H3K9ac and H3K18ac data are from Ferrari et al. 2012 [26]. Replication sequencing data are from Encode Consortium (www.encodeproject.org). Other chromatin feature data were downloaded from NCBI epigenome roadmap project (www.ncbi.nlm.nih.gov/geo/roadmap/epigenomics/).

We also compared VGI frequencies for regions in different cytogenic categories. The mean VGI frequency decreases with increasing Giemsa staining levels (for bins in categories ranging from “gneg” to the “gpos100” cytogenetic categories) (Figure 3c). Centromeric stained regions (categorized as “acen” in Figure 3c) are heterochromatic and have the lowest average VGI frequencies. This observation also suggests that the adenovirus genome interacts less frequently with heterochromatin and much more so with active euchromatin regions.

We further analyzed the viral-host genome contacts in the context of data from DNase I hypersensitivity assays (Figure S2) and ChIP-seq assays of various histone modifications (Figure 3d). These analyses indicated that viral DNA preferentially interacts with active open chromatin regions. For instance, the 222 VGI enriched regions at 6 hours p.i. show a similar histone modification profile as those in compartment A regions (denoted as VGI and A, respectively, in Figure 3d), whose histone modification markers typically are associated with actively transcribed chromatin regions such as H3K4me1, H3Kme2, H3K27ac, H3K79me1 and H3K79me2 [35,36]. Histone modifications H3K27ac and H3K4me1 were previously shown to be defining features of active enhancer regions [36]. Even the 74 VGI regions classified as being part of the B compartment (denoted as VGI_B in Figure 3d) showed higher fold change for histone modifications associated with active open chromatin (i.e. H3K27ac, H3K4me1, H3K4me2, etc.) compared with the chromatin regions in the B compartment (denoted as B in Figure 3d). In addition, we observed a strong negative correlation (Pearson’s correlation coefficient = −0.39 in Figure S2) and a depleted fold change (fold change = 0.5 in Figure 3d) between the VGI frequency profile and the occurrence profile of histone modification H3K9me3, which is a marker for heterochromatin regions [37,38]. These observations suggest that viral DNA-host genome interactions occur preferentially in active open chromatin regions. This enrichment could be the result of preferential binding of the viral genome to active chromatin regions, an increased viral DNA replication in these active regions, active exclusion from heterochromatin, or combinations of these mechanisms. Since our analyses were based on data from 6 hours p.i. when no significant viral replication is taking place, and viral DNA interacts with specific regions of host euchromatin, we favor the interpretation that the viral genome preferentially interacts with open active chromatin regions.

Interestingly, histone marker H3K27me3, which is associated with Polycomb-group silencing [35,23], is strongly enriched in the VGI genomic regions (VGI, VGI_A, and VGI_B in Figure 3d), even though it antagonizes H3K27ac, a histone modification enriched in compartment A but not in compartment B regions and associated with gene activity [39]. Given their opposite transcriptional activities, it is intriguing that both H3K27ac and H3K27me3 markers are enriched at VGI regions (especially in VGI_A regions). Both H3K27ac and H3K27me3 are involved in transcription regulation of genes responsible for cell differentiation [36]. Whether their enrichment in VGI regions is mechanistically linked to viral induced transformation remains an interesting open question. Nevertheless, these analyses suggest that viral-host genome interactions may define unique chromatin states, which appear to share many features of the open euchromatin but also show distinct histone marker profile from those associated with activated and repressed genes.

### Relations between viral DNA contacts and DNA replication

Viral DNA replication occurs within defined regions of the host cell’s nucleus [40,41]. Figure 3e shows that compartment A regions, particularly the 148 VGI’s that are part of compartment A (denoted as VGI_A), are early replicated. Even the 74 VGI regions classified as being part of the B compartment showed substantially earlier replication timing compared with other regions in the B compartment, which generally replicate at later stages (Figure 3e; VGI_B regions showed highest replication signals in the mid S phase, while compartment B regions’ signals generally enrich towards the G2 phase.). These observations suggest that the adenovirus genome prefers to interact with the host genome’s early replication regions.

Infection with the *dl1500* adenovirus will drive arrested fibroblast cells into S phase [25,21]. To see if the host genome replication driven by *dl1500* adenovirus infection spatially co-localizes with positions of viral DNA attachments, we performed BrdU-seq experiments in 24 hours p.i. IMR90 cells following experimental procedures as previously described [42]. IMR90 cells were treated with BrdU at 23.5 hours post *dl1500* adenovirus infection. Cells were harvested 30 min later and BrdU incorporated DNA were immunoprecipitated and purified for high-throughput sequencing. Sequencing reads were aligned to both the hg19 and Ad5 reference genomes. Among the uniquely aligned reads, 95% aligned to the Ad5 reference genome, indicating that the majority of DNA replication is from the virus genome (Figure 4a). The 5% BrdU-seq reads aligned to hg19, a total number of 1,009,498 reads, were indexed to 500-kb resolution and compared with the VGI contact profiles at 24 hours pi (Figure 4b). Among the 731 VGI enriched chromatin bins at 24 hours p.i., about 40% overlapped with BrdU-seq signals. Of the remaining bins without virus-genome interactions, only ~20% showed host genome replication signals (Figure 4c). However, since genomic regions with BrdU-seq signal are also much more early replicated than the other regions (Figure S3; p-value less than 10^−16^), we cannot determine if host genome replication at 24 hours p.i. is either due to its intrinsic early replication timing or driven by viral activities. Based on the BrdU-seq data, we observed that the majority of the DNA replication activities happened to the virus genome, which is likely the result of high MOI value of virus infection [21]. Therefore, the fact that more genomic regions are identified with VGI enrichment in 24 vs. in 6 hours p.i. (731 vs. 222) could be due to extra viral genome load at the later time point.

**Figure 4.**
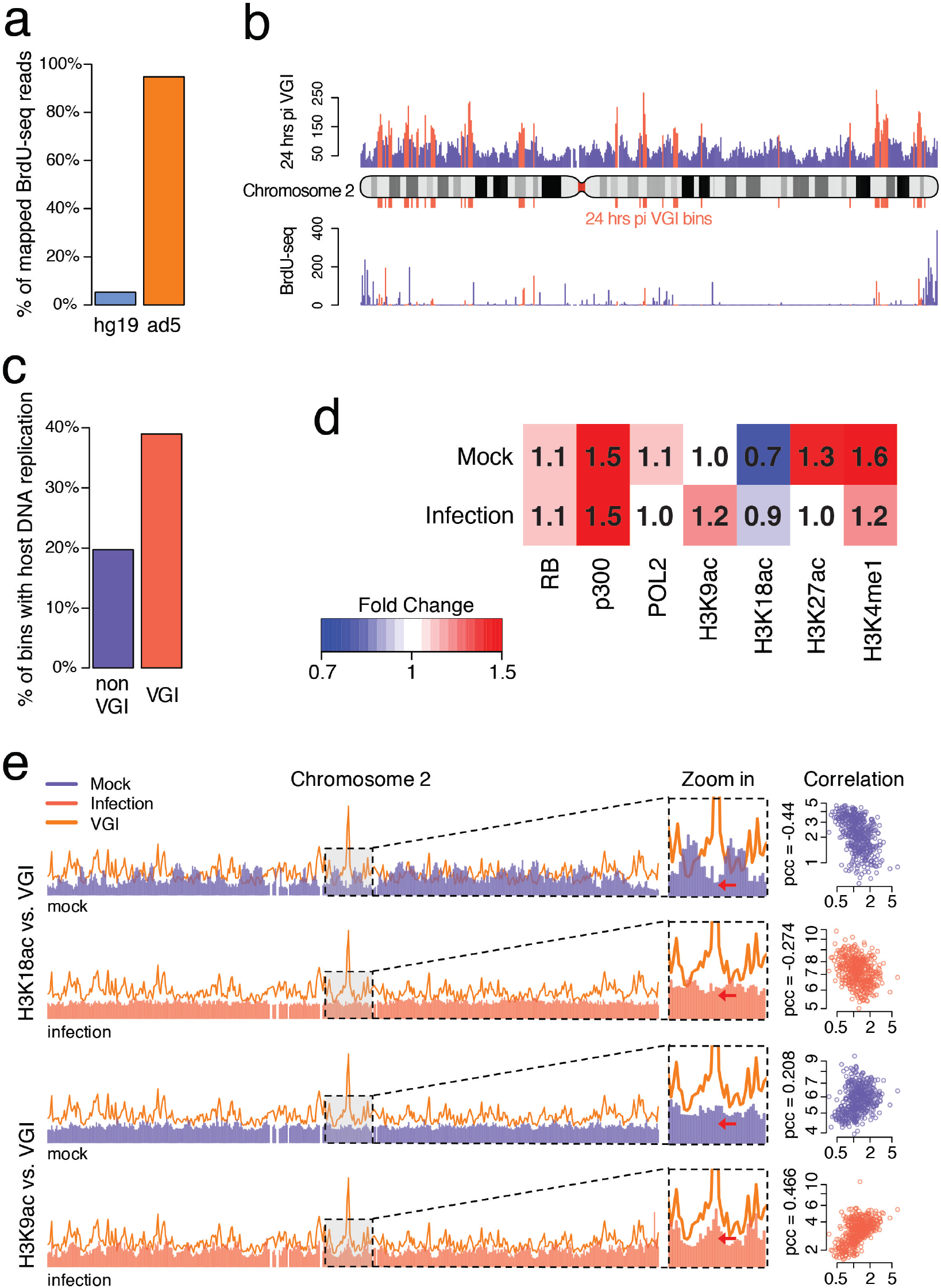
Virus and host genome replication. Epigenetic remodeling at VGI enriched bins. a) Fraction of BrdU-seq reads alignment to either hg19 or Ad5 reference genomes. b) VGI contact frequency profile (upper panel) and BrdU-seq frequencies (lower panel) at 24 hours pi for chromosome 2. VGI contact frequencies and BrdU-seq reads counts of VGI enriched bins are colored in red while all data of all other bins are colored in blue. All the profiles are based on 500kb resolution. c) Percentage of VGI enriched bins with host genome replication identified by BrdU-seq (in red). For the other bins, the percentage is colored in blue. All data shown for 24 hours pi. d) Heatmap of different ChIP-seq data before and after 24 hours of adenovirus *dl1500* infection at the 24 hours pi VGI enriched host genome regions. Strengths of signals for the heatmap entries are represented by fold change in VGI enriched regions compared to the genome-wide average signals. Sources of data are from Ferrari et al. 2012 [26] and Ferrari et al. 2014 [27]. e) 6 hours pi VGI contact frequency distribution on Chromosome 2. H3K18ac and H3K9ac distributions for mock-infection and post-infection samples. Zoom in regions show alterations of different histone acetylations after infection. Correlations between primary VGI contact profile and either H3K18ac or H3K9ac distribution in either mock-infection or post-infection sample.

### Remodeling of epigenetic features in the VGI enriched host genomic regions

Since small e1a alters global patterns of specific histone modifications in the host genome [24-26], we compared ChIP-seq data of various histone modifications before and after adenovirus *dl1500* infection. We found that VGI enriched genomic regions undergo dramatic changes in their epigenetic chromatin features upon viral infection (Figure 4d). The host lysine acetylase p300 binds to VGI enriched regions after infection. p300 interacts with the retinoblastoma (RB) proteins, which together with e1a (the p300-e1a-RB complex) function in repressing selected host genes involved in anti-viral defense [27]. Reduced fold changes of H3K27ac and H3K4me1 were observed in the VGI enriched genomic regions after infection (Figure 4d), indicating that VGI regions coincide with locations of host epigenome remodeling. In addition, the histone modification profile of H3K18ac before infection is strongly anti-correlated (Pearson’s correlation coefficient = −0.44) with the VGI frequencies at early infection (Figure 3d, 4d, S2). Upon viral infection small e1a causes a substantial (~70%) reduction in cellular levels of H3K18ac and specifically, the elimination of essentially all H3K18ac ChIP-seq peaks at promoters and intergenic regions of genes related to fibroblast functions [25,26]. Meanwhile, the small e1a also induces the appearance of new H3K18ac peaks at promoters of highly induced genes associated with cell cycle control and at new putative enhancers [26]. It is interesting to see that the viral genome preferentially interacts with the host genome at positions deprived of H3K18ac at early stages of infection. And after 24 hours of infection, H3K18ac is enhanced at such VGI genomic regions in the presence of RB and p300 (Figure 4d, 4e). These observations suggest that acetylation of H3K18 [25], may be related to the binding of the adenovirus genome or shows similar preference for chromatin regions associated with viral genome interactions. Besides H3K18ac, enrichment is also found for H3K9ac in the VGI enriched genomic regions because of the general colocalization of these two histone acetylation marks [26,43].

The above observations suggest that virus genome attachments to the host genomic regions coincide and may even potentially induce epigenetic changes during adenovirus infection. The mechanisms by which the binding of the virus genome may be related causally or consequentially to such epigenetic changes are unclear. But our studies show non-random viral-host genome interactions and raise an interesting question for future studies on the role of VGI in host epigenome remodeling, host gene expression, viral genome replication and the underlying mechanisms of oncogenic transformation caused by adenovirus infection.

## Conclusions

In this study, we have analyzed the global physical interactions between the adenovirus genome and host fibroblast genome using tethered chromosome conformation capture. Our analyses indicate that virus-host genome interactions occur at specific genomic locations. By mapping VGIs in the host genome at different stages of adenovirus infection, we find that the initial pattern of VGI locations is largely maintained throughout the infection process and at later stage of infection the distribution of VGIs expands to additional host genomic regions. VGIs preferentially occur in the active chromatin subcompartment in the host genome, which could potentially facilitate viral gene expression and DNA replication. While the host genomic regions targeted preferentially by virus interactions share many epigenetic features of activated genes, they also possess many unique histone markers, suggesting that virus attachment sites on the host genome constitute a unique class of chromatin states. Our analyses also suggest that VGI could have substantial impact on the activities of the host genome. Adenovirus-induced epigenetic remodeling of the host genome appears to be enhanced in genomic regions enriched with VGIs. Likewise, the binding of regulatory protein machineries (the p300 and RB complex) is increased at chromatin regions with enriched VGIs. It is conceivable that VGIs may be facilitating these changes or that these epigenetic changes facilitate VGIs to achieve a favorable outcome for the virus. Further experiments with higher resolutions are needed to validate and expand on these findings, to reveal more detailed structural features of the host chromatin regions where virus genome preferentially bind, and to elucidate the role of virus genome in regulating host genome activities.

## Methods

### Cell culture and TCC experiments

IMR90 cells were grown under standard culturing conditions (DMEM, 10% FBS, 1X penicillin/streptomycin, 5% CO_2_, and 37 °C). Once the cultures reached 100% confluency, they were maintained for another 12 hours to make sure cells were being arrested. Cells were infected with small e1a adenovirus [26] (MOI=200) in regular media while FBS was reduced to 2% and maintained at 37 °C with 5% CO_2_. Mock infection cells received the new media with no virus. At each stage of 6/24 hours post-infection (pi) and 6/24 hours mock-infection (mi), old media was aspirated out and 20 millions cells were treated with fresh medium (DMEM, 2% FBS, and penicillin/streptomycin) and 1% of formaldehyde. TCC experiments were performed as previously described [15].

### Preprocessing TCC sequencing output

Sequencing libraries prepared from TCC experiments have unique features because of a number of special enzyme treatments. In this section, preprocessing of the raw sequencing reads prior to alignment is described.

#### Adjusting for sequencing lariats

TCC experiments use exonuclease III to remove the free DNA ends’ biotin before pulling down of DNA ligation junctions. This step may produce a significant amount of single strand DNA, which can potentially form hairpin structures during preparation of the sequencing library. When performing DNA end repair, 5 prime overhang of hairpin structure will be filled in, which cause the initial sequence of read 2 to be identical to what from the read 1 and eventually lead to significant amount of alignment failures. We name this type of errors as “sequencing lariats” which is unique for TCC experiments. To attenuate the effect of these errors, we run a program to search for identical initial sequences between read 1 and read 2 and trim the identical sequence from read 2 before performing the alignment. This trimming step significantly increased the percentage of alignable reads.

#### Filtering of ligation junctions

Informative sequencing results are the ligation products of different DNA fragments resulted from restriction enzyme digestion. Depending on the sites of DNA shearing, sequencing reads may surpass the ligation junctions of certain DNA fragments. These reads may be unable to align to the genome because of the chimeric ligations. In order to improve the alignment efficiency, we run a program to scan for HindIII ligation junction (expected to be “AAGCTAGCTT” after end filling-in and blunt-end ligation) and allow one base of mutation or deletion considering restriction enzyme star effect) and removed all bases after the 3 prime midpoint of the junction. This filtering step also led to bigger portion of successfully aligned reads.

Raw sequencing reads after the previous two steps of filtering are aligned to hg19 and Adenovirus 5 reference genome by bowtie-1.0.0 with a maximum of three mismatches allowed.

#### Removing non-informative pairs

Two types of sequencing read pairs that do not contain any information about the spatial organization of the genomes and viral-host genome interactions can be readily identified. The first type consists of pairs that are a result of PCR amplification of a single DNA molecule (PCR duplication). These pairs can make a contact appear more frequent when it has only been amplified more efficiently in the PCR. A group of read pairs that are a result of PCR duplication align to identical positions on both ends. All but one pair in such groups were removed from the catalogue.

The second type consists of pairs that originate from DNA molecules that do not include a ligation junction, yet they bind to the streptavidin-coated beads either non-specifically or as a result of incomplete exonuclease action in removing terminal biotins. After sequencing, these molecules result in pairs that align just 300-700 bp apart to opposite strands of the reference sequence of the genome (the size range depends on the size range used during gel-extraction). All such pairs were removed from the catalogue of each dataset.

### Splitting test for filtering low coverage bins of TCC observed contact (OC) matrix and choosing optimal resolution to visualize VGI contact profile

Difference in chromatin accessibility of genomic locations leads to difference in sequencing depth and coverage, which requires special considerations in most of the sequencing based genomic analyses [44]. For TCC and Hi-C studies, because of low-coverage in 2D contact matrices, sequencing depth is one of the key considerations. Values of low coverage matrix entries won’t converge during iterative normalization [33], which leads to spurious contact frequencies (Figure S4).

#### Splitting test for TCC OC matrix

To define matrix bins of low coverage, we performed a “splitting test” on the TCC OC matrix, which will divide the data matrix into two matrices by randomly dividing the unique read counts per entry bin (with value n) following a binomial distribution (x ~ ***binom***(*n, p*=0.5)). The pair of matrices will be used to test if contact profiles at the given bin resolution are reproducible. Several pairs of matrices will be generated by independent splitting test of the original data. Splitting tests were performed on matrices from all of the four TCC datasets.

Intrachromosomal contacts are not random and maintain specific patterns in interphase cells [12,15,45]. Therefore, the Pearson’s correlation between the corresponding pairs of intrachromosomal contact profiles in both split matrices should be close to +1 if the bin size is adequately chosen with respect to the available sequencing coverage. Splitting tests showed that at 500kb bin resolution, the pair-wise Pearson’s correlation converges to +1 at 1%~2% percentiles. Bins with correlations in the lower 1%~2% percentiles will be removed from the matrices.

#### Splitting test for VGI contact vector

To determine the optimal resolution to present the viral-host genome interactions (VGIs), we performed the same binomial splitting for the VGI contact vector entries and compared the pairwise correlations at a series of different resolutions. Each VGI contact vector was split independently 20 times to test the consistency between the splitting tests. Similar as interchromosomal interactions, VGIs are more uniformly distributed than intrachromosomal interactions. Therefore, pairwise Pearson’s correlation of VGI splitting tests do not converge to 1 at any resolutions in either one of the two datasets (6 hours pi and 24 hours pi). So here we choose a reasonable resolution of 500kb, when the pairwise correlations are generally high and starting to converge (Figure S1).

### Bias correction of TCC datasets

#### Iterative correction of TCC OC matrix

TCC OC matrices of the four datasets are normalized through the iterative correction (ICE) method to remove experimental biases [33]. As described in the ICE method, the diagonal of each OC matrix was removed. Also all intrachromosomal interactions with genomic distances less than 20kb were removed, which removes potential self-looping contacts and dangling ends artifacts between consecutive bins. OC matrix bins with low sequencing coverage (as defined by splitting tests) were removed resulting in reduced matrix dimension. At 500kb resolution, each TCC OC matrix was defined as a *K*×*K* matrix, ***C*** = (*c_ij_*)_*K*×*K*_, in which the entry *c_ij_* is equal to the number of observed CIs between bins *i* and *j*, where *K* is the matrix dimension after removing low coverage bins and *c_ij_* = 0 for *i* = *j*. To obtain the contact frequency (CF) matrix ***F_n_*** = (*f_ij,n_*)_*K*×*K*_ through ICE method, the matrix was normalized for *n* iterations as follows:

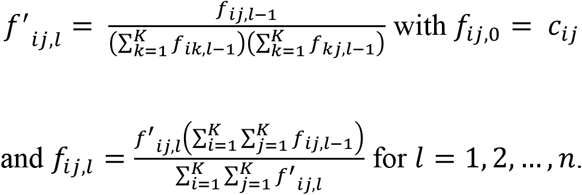

Therefore, the bias vector is ***V_n_*** = (*v_n,i_*)_*K*_ where 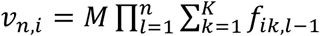 with the scaling factor 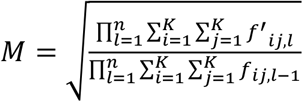 so that 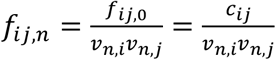. OC matrix for each dataset was normalized for 10 iterations when the matrix entries started to converge.

#### Normalization of VGI

To obtain contact frequency of VGI vector, ***A*** = (*a_i_*)_*K*_, at 500kb resolution, the observed VGI count vector ***O*** = (*o_i_*)_*K*_ was normalized by its corresponding bias vector ***V_n_*** so that 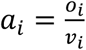. The bias vector ***V_n_*** was also scaled to 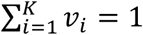 to serve as the probability vector in identifying VGI enriched regions on the host genome.

### Identification of VGI enriched genomic regions by hypothesis testing

A multinomial null model assumption was used for the VGI contact distribution on genomic locations with *Mult_k_(n, **p**)*, for which *k* is the number of binning windows (*k*=5,644), *n* is the total number of observed VGIs for the 6 hours pi and 24 hours pi sample, and ***p*** is the binning windows’ probability vector (*p_1_, p_2_*, …, *p_k_*), which is proportional to the OC matrix bias vector generated by the ICE normalization. Bins with significantly higher number of VGIs than expected by the random null model distribution were defined as VGI enriched with a false discovery rate (FDR) of less than 10^−4^. The FDR was calculated by applying the same statistics to two randomly generated null models given different p-value thresholds.

## Supplementary Materials

**Figure s1.**
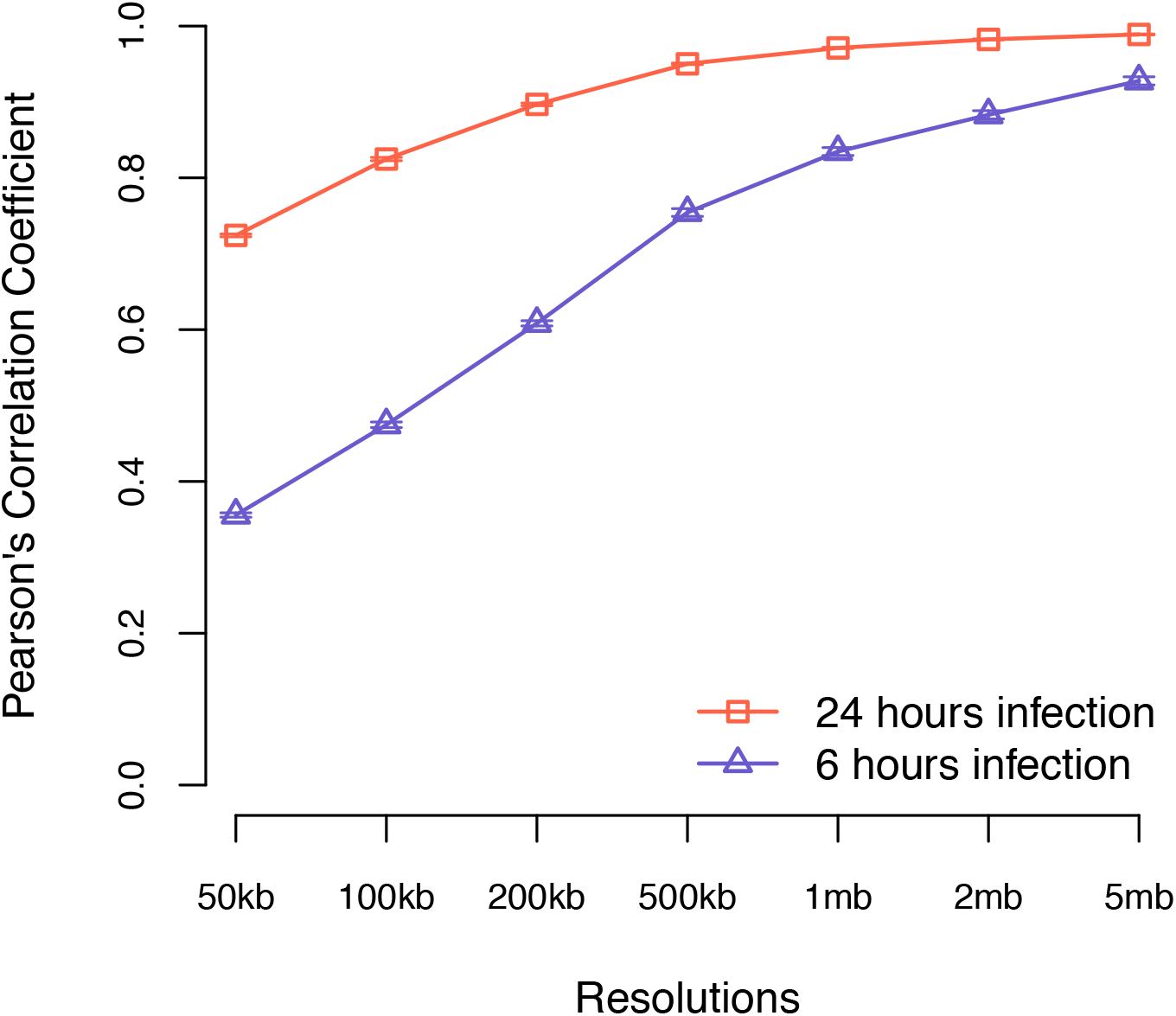
Splitting test of viral-host genome interactions at different bin resolutions. VGIs from 6 hours pi and 24 hours pi datasets were indexed at different resolutions ranging from 50kb to 5mb. VGI contact vectors with different resolutions were randomly split using a binomial distribution for 20 times (Methods) and Pearson’s correlations between two split vectors were calculated for each splitting event (Methods). The error bar at different resolutions represents the standard deviation of the pairwise correlations among the 20 times of splitting.

**Figure s2.**
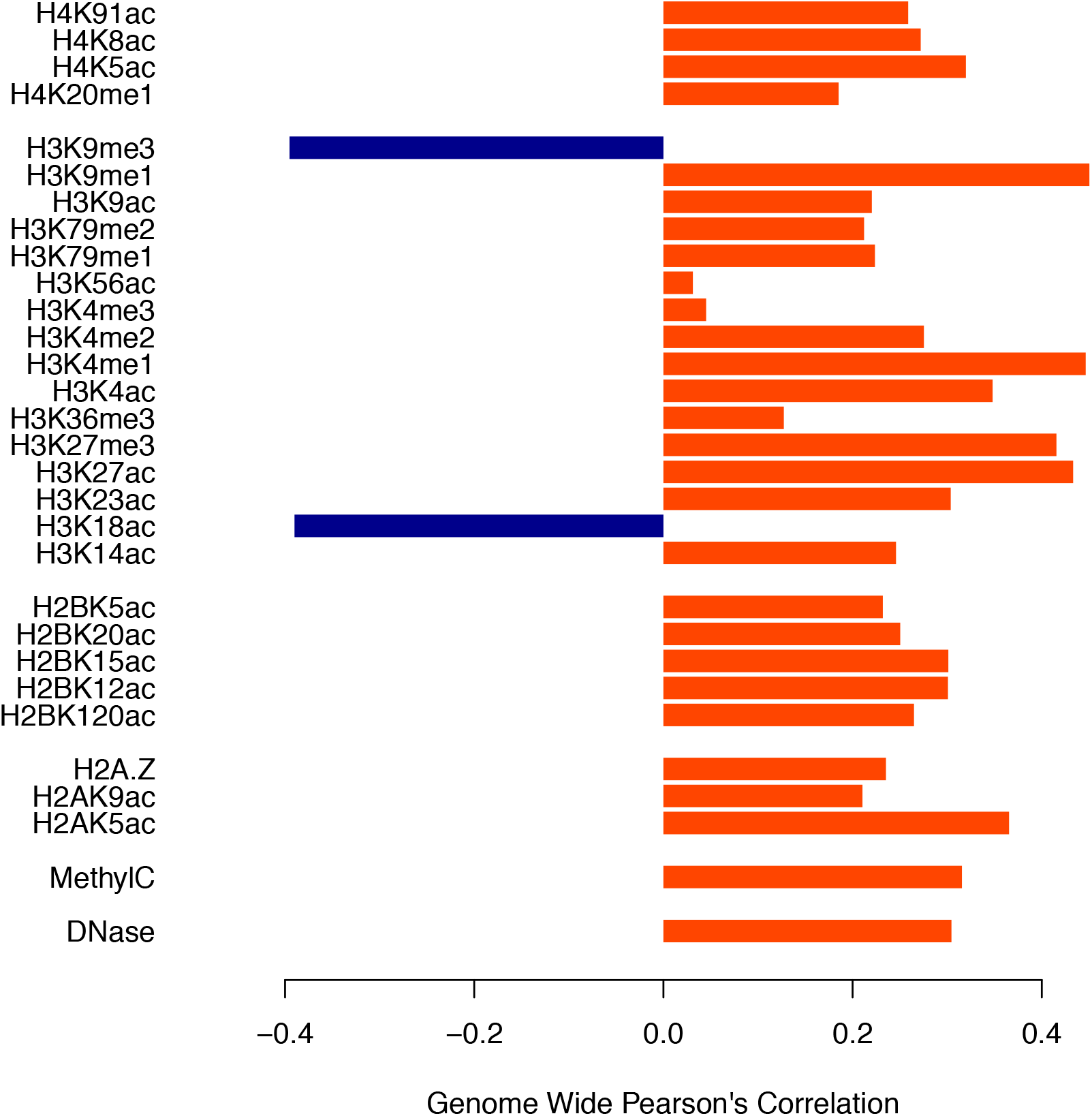
Genome wide Pearson’s correlations between 6 hours VGI contact frequency profile and different chromatin feature profiles. H3K9ac and H3K18ac Chip-Seq data were downloaded from the study Ferrari, et al [26]. Other chromatin feature data was downloaded from NCBI epigenome roadmap project.

**Figure s3.**
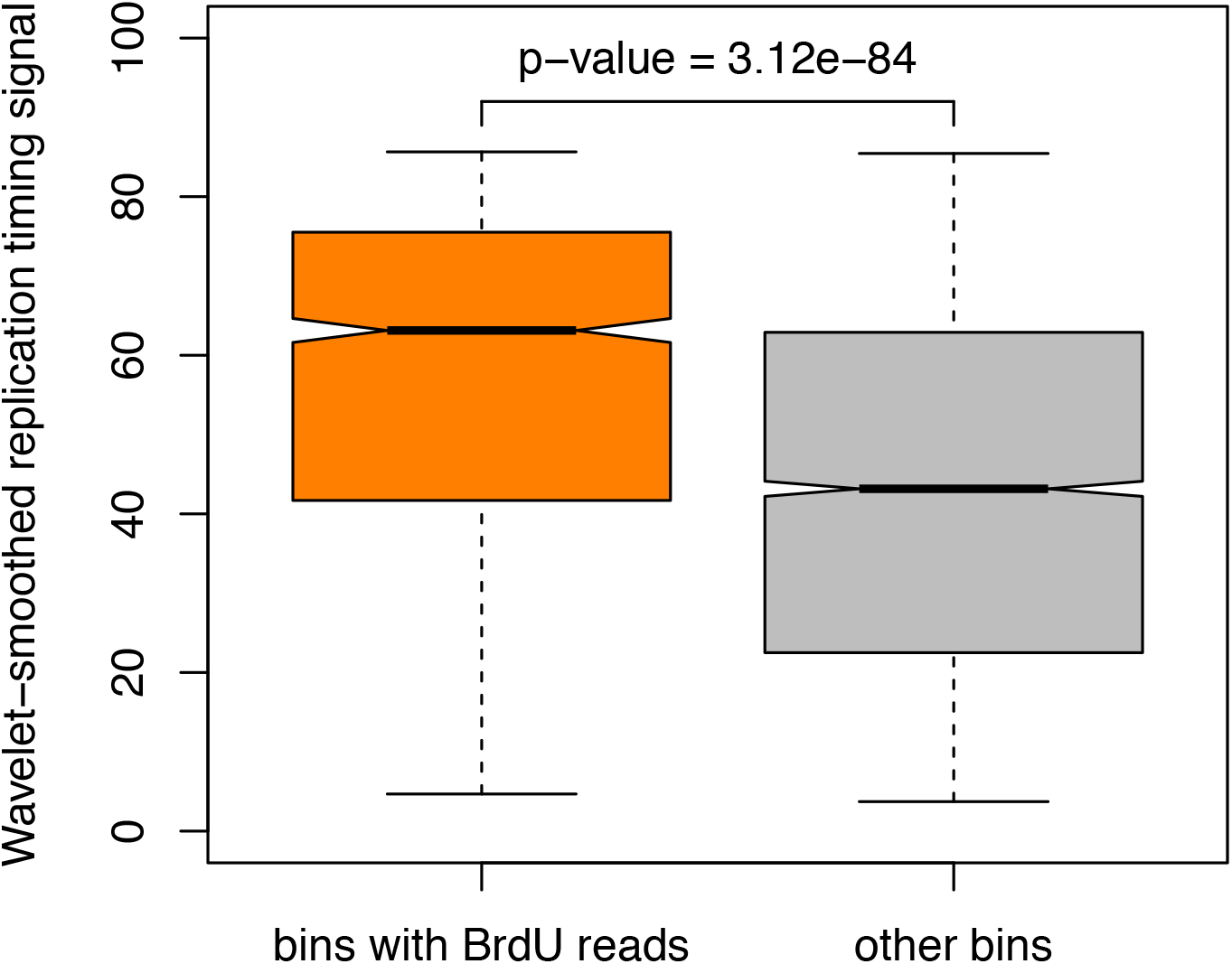
Comparison of replication timing signal between genomic regions with BrdU-seq reads at 24 hours pi versus regions without BrdU-seq reads. The Wavelet-smoothed replication timing signal are retrieved from the ENCODE project consortium database.

**Figure s4.**
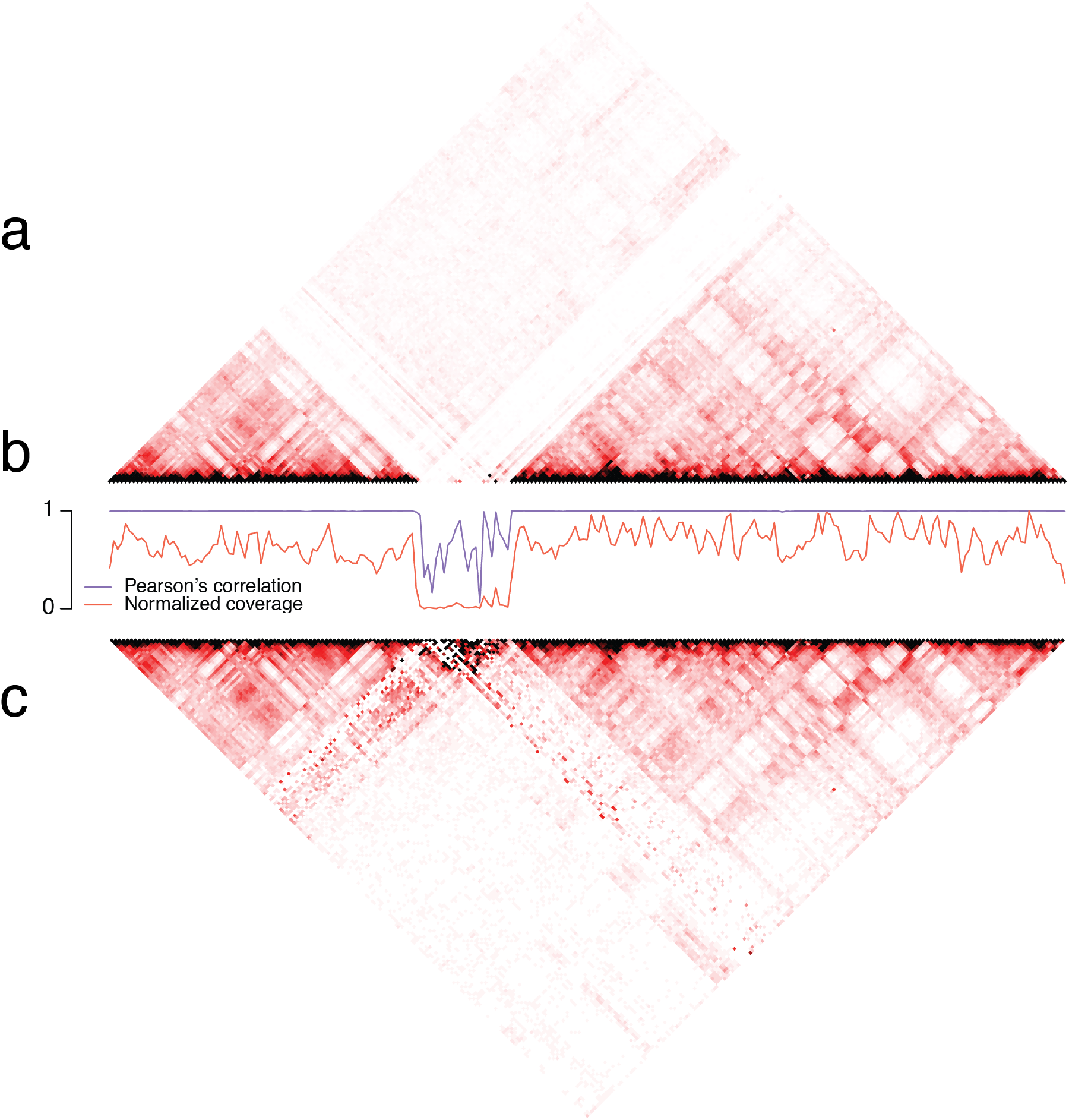
Splitting test of chromosomal TCC contact matrix. a) Chromosome 2 observed contact (OC) matrix b) Splitting test: Purple line shows the pair-wise Pearrson’s correlations between split matrices for different bins. Orange line shows the normalized read coverage for different bins. c) Spurious contact frequencies appear at low coverage bins of contact frequency (CF) matrix after iterative correction.

**Table s1.**
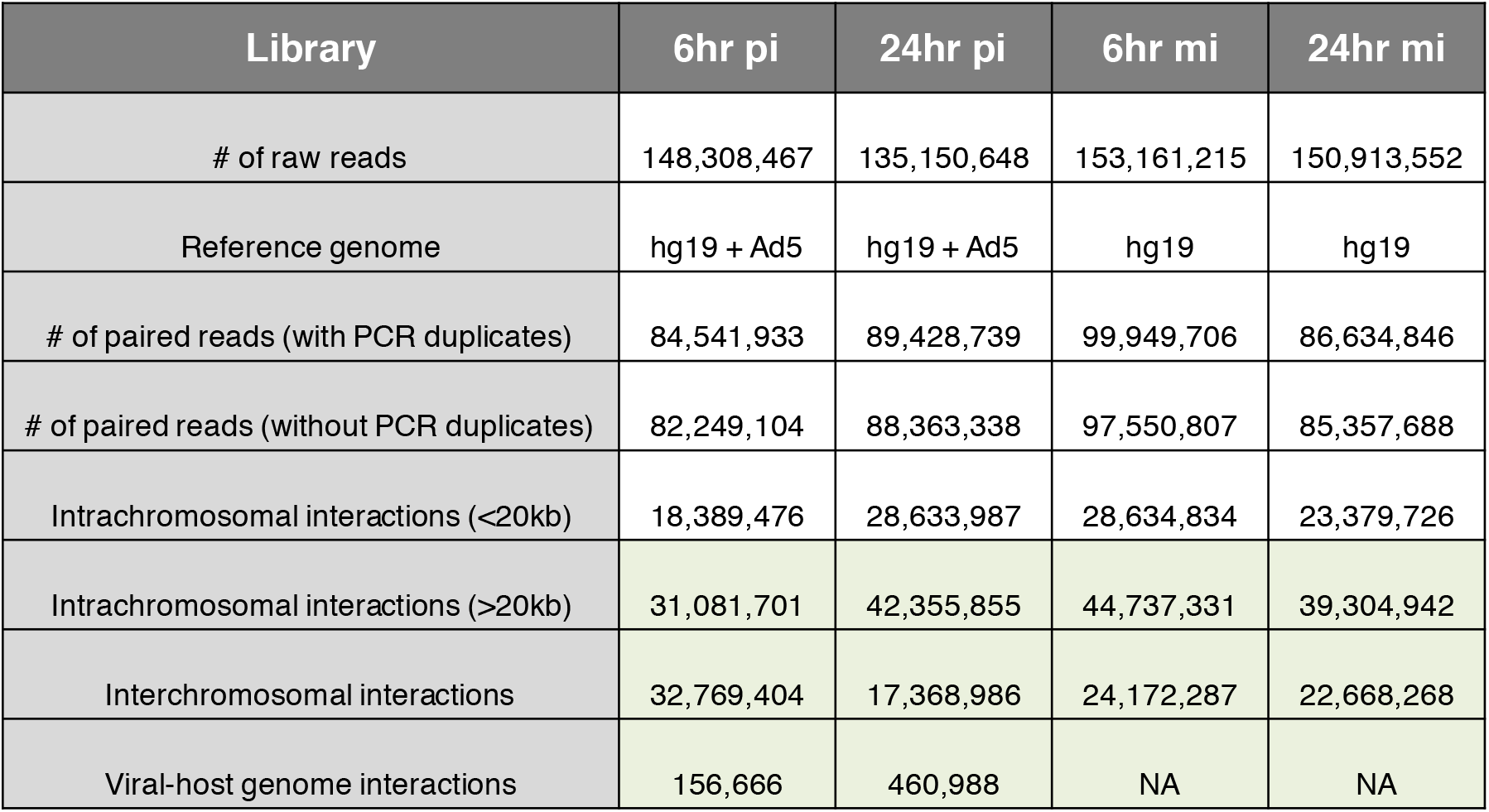
Summary of the numbers of sequencing reads at each step of TCC data processing.

**Table s2.** Summary of VGI distributions between compartment A and B regions for different chromosomes.

| Chromosome | Compartment with higher VGI | P-value |
| --- | --- | --- |
| chr1 | A | 0.00491 |
| chr2 | A | 1.37e-12 |
| chr3 | A | 9.35e-09 |
| chr4 | A | 2.5e-19 |
| chr5 | A | 3.02e-15 |
| chr6 | A | 2.84e-09 |
| chr7 | A | 1.86e-06 |
| chr8 | A | 5.02e-11 |
| chr9 | A | 1.58e-05 |
| chr10 | A | 8.39e-05 |
| chr11 | A | 0.00109 |
| chr12 | A | 0.0134 |
| chr13 | A | 3.98e-06 |
| chr14 | A | 0.000873 |
| chr15 | A | 0.0667 |
| chr16 | B | 0.764 |
| chr17 | B | 8.14e-10 |
| chr18 | A | 1.29e-06 |
| chr19 | B | 0.972 |
| chr20 | A | 0.369 |
| chr21 | A | 0.121 |
| chr22 | B | 0.52 |
| chrX | A | 7.63e-29 |

## References

1. Schmid M, Speiseder T, Dobner T, Gonzalez RA. DNA Virus Replication Compartments. J. Virol. 2014;88:1404–20.

2. Rivas H, Schmaling S, Gaglia M. Shutoff of Host Gene Expression in Influenza A Virus and Herpesviruses: Similar Mechanisms and Common Themes. Viruses. 2016;8:102.

3. Slinger E, Langemeijer E, Siderius M, Vischer HF, Smit MJ. Herpesvirus-encoded GPCRs rewire cellular signaling. Mol. Cell. Endocrinol. 2011;331:179–84.

4. Knipe DM, Cliffe A. Chromatin control of herpes simplex virus lytic and latent infection. Nat. Rev. Microbiol. 2008;6:211–21.

5. Lieberman PM. Chromatin organization and virus gene expression. J. Cell. Physiol. 2008. p. 295–302.

6. Maul GG. Nuclear domain 10, the site of DNA virus transcription and replication. BioEssays. 1998. p. 660–7.

7. Ishov AM, Maul GG. The periphery of nuclear domain 10 (ND10) as site of DNA virus deposition. J. Cell Biol. 1996;134:815–26.

8. Cremer T, Cremer C. Chromosome territories, nuclear architecture and gene regulation in mammalian cells. Nat Rev Genet. 2001;2:292–301.

9. Misteli T. Beyond the Sequence: Cellular Organization of Genome Function. Cell. 2007. p. 787–800.

10. Bickmore W a. The spatial organization of the human genome. Annu. Rev. Genomics Hum. Genet. 2013;14:67–84.

11. Dekker J, Rippe K, Dekker M, Kleckner N. Capturing chromosome conformation. Science. 2002;295:1306–11.

12. Lieberman-Aiden E, van Berkum NL, Williams L, Imakaev M, Ragoczy T, Telling A, et al. Comprehensive mapping of long-range interactions reveals folding principles of the human genome. Science. 2009;326:289–93.

13. Rao SSPSP, Huntley MHH, Durand NCC, Stamenova EKK, Bochkov IDD, Robinson JTT, et al. A 3D Map of the Human Genome at Kilobase Resolution Reveals Principles of Chromatin Looping. Cell. 2014;159:1665–80.

14. Dixon JR, Selvaraj S, Yue F, Kim A, Li Y, Shen Y, et al. Topological domains in mammalian genomes identified by analysis of chromatin interactions. Nature. 2012;485:376–80.

15. Kalhor R, Tjong H, Jayathilaka N, Alber F, Chen L. Genome architectures revealed by tethered chromosome conformation capture and population-based modeling. Nat. Biotechnol.. 2012;30:90–8.

16. Dekker J. The three “C” s of chromosome conformation capture: controls, controls, controls. Nat. Methods. 2006;3:17–21.

17. Fullwood MJ, Liu MH, Pan YF, Liu J, Xu H, Mohamed Y Bin, et al. An oestrogen-receptor-alpha-bound human chromatin interactome. Nature. 2009;462:58–64.

18. Gavrilov AA, Gushchanskaya ES, Strelkova O, Zhironkina O, Kireev II, Iarovaia O V., et al. Disclosure of a structural milieu for the proximity ligation reveals the elusive nature of an active chromatin hub. Nucleic Acids Res. 2013;41:3563–75.

19. Nagano T, Lubling Y, Stevens TJ, Schoenfelder S, Yaffe E, Dean W, et al. Single-cell Hi-C reveals cell-to-cell variability in chromosome structure. Nature. 2013;502:59–64.

20. Trentin JJ, Yabe Y, Taylor G. The quest for human cancer viruses. Science. 1962;137:835–41.

21. Berk AJ. Recent lessons in gene expression, cell cycle control, and cell biology from adenovirus. Oncogene. 2005;24:7673–85.

22. Davison AJ, Benko M, Harrach B. Genetic content and evolution of adenoviruses. J. Gen. Virol. 2003. p.2895–908.

23. Cao R. Role of Histone H3 Lysine 27 Methylation in Polycomb-Group Silencing. Science. 2002;298:1039–43.

24. Ferrari R, Pellegrini M, Horwitz G a, Xie W, Berk AJ, Kurdistani SK. Epigenetic reprogramming by adenovirus e1a. Science. 2008;321:1086–8.

25. Horwitz GA, Zhang K, McBrian MA, Grunstein M, Kurdistani SK, Berk AJ. Adenovirus Small e1a Alters Global Patterns of Histone Modification. Science. 2008;321:1084–5.

26. Ferrari R, Su T, Li B, Bonora G, Oberai A, Chan Y, et al. Reorganization of the host epigenome by a viral oncogene. Genome Res. 2012;22:1212–21.

27. Ferrari R, Gou D, Jawdekar G, Johnson SA, Nava M, Su T, et al. Adenovirus Small E1A Employs the Lysine Acetylases p300 / CBP and Tumor Suppressor Rb to Repress Select Host Genes and Promote Productive Virus Infection. Cell Host Microbe. 2014;16:663–76.

28. Montell C, Fisher EF, Caruthers MH, Berk AJ. Resolving the functions of overlapping viral genes by site-specific mutagenesis at a mRNA splice site. Nature. 1982;295:380–4.

29. Liu X, Marmorstein R. Structure of the retinoblastoma protein bound to adenovirus E1A reveals the molecular basis for viral oncoprotein inactivation of a tumor suppressor. Genes Dev. 2007;21:2711–6.

30. Yaffe E, Tanay A. Probabilistic modeling of Hi-C contact maps eliminates systematic biases to characterize global chromosomal architecture. Nat. Genet. 2011;43:1059–65.

31. Hu M, Deng K, Selvaraj S, Qin Z, Ren B, Liu JS. HiCNorm: Removing biases in Hi-C data via Poisson regression. Bioinformatics. 2012;28:3131–3.

32. Hahn S, Kim D. Identifying and Reducing Systematic Errors in Chromosome Conformation Capture Data. PLoS One. 2015;10:e0146007.

33. Imakaev M, Fudenberg G, McCord RP, Naumova N, Goloborodko A, Lajoie BR, et al. Iterative correction of Hi-C data reveals hallmarks of chromosome organization. - Supplement. Nat. Methods. 2012;9:999–1003.

34. Frey MR, Matera a G. Coiled bodies contain U7 small nuclear RNA and associate with specific DNA sequences in interphase human cells. Proc. Natl. Acad. Sci. U. S. A. 1995;92:5915–9.

35. Martin C, Zhang Y. The diverse functions of histone lysine methylation. Nat. Rev. Mol. Cell Biol. 2005;6:838–49.

36. Creyghton MP, Cheng AW, Welstead GG, Kooistra T, Carey BW, Steine EJ, et al. Histone H3K27ac separates active from poised enhancers and predicts developmental state. Proc. Natl. Acad. Sci. 2010;107:21931–6.

37. Nakayama J, Rice JC, Strahl BD, Allis CD, Grewal SI. Role of histone H3 lysine 9 methylation in epigenetic control of heterochromatin assembly. Science. 2001;292:110–3.

38. Rea S, Eisenhaber F, O’Carroll D, Strahl BD, Sun ZW, Schmid M, et al. Regulation of chromatin structure by site-specific histone H3 methyltransferases. Nature. 2000;406:593–9.

39. Tie F, Banerjee R, Stratton C a, Prasad-Sinha J, Stepanik V, Zlobin A, et al. CBP-mediated acetylation of histone H3 lysine 27 antagonizes Drosophila Polycomb silencing. Development. 2009;136:3131–41.

40. de Bruyn Kops a, Knipe DM. Preexisting nuclear architecture defines the intranuclear location of herpesvirus DNA replication structures. J. Virol. 1994;68:3512–26.

41. Wang IH, Suomalainen M, Andriasyan V, Kilcher S, Mercer J, Neef A, et al. Tracking viral genomes in host cells at single-molecule resolution. Cell Host Microbe. 2013;14:468–80.

42. Li B, Su T, Ferrari R, Li JY, Kurdistani SK. A unique epigenetic signature is associated with active DNA replication loci in human embryonic stem cells. Epigenetics. 2014;9:257–67.

43. Jin Q, Yu L-R, Wang L, Zhang Z, Kasper LH, Lee J-E, et al. Distinct roles of GCN5/PCAF-mediated H3K9ac and CBP/p300-mediated H3K18/27ac in nuclear receptor transactivation. EMBO J. 2011;30:249–62.

44. Sims D, Sudbery I, Ilott NE, Heger A, Ponting CP. Sequencing depth and coverage: key considerations in genomic analyses. Nat. Rev. Genet. 2014;15:121–32.

45. Naumova N, Imakaev M, Fudenberg G, Zhan Y, Lajoie BR, Mirny L a, et al. Organization of the mitotic chromosome. Science. 2013;342:948–53.

